# Computational Fluid Dynamics Modeling of Aerosol Particle Transport through Lung Airway Mucosa

**DOI:** 10.1101/2021.10.18.464809

**Authors:** Blake A. Bartlett, Yu Feng, Catherine A. Fromen, Ashlee N. Ford Versypt

## Abstract

Delivery of aerosols to the lung has great potential for the treatment of various lung diseases. However, the lungs are coated by a protective mucus layer whose complex properties make this form of delivery difficult. Mucus is a non-Newtonian fluid and is cleared from the lungs over time by ciliated cells. Further, its gel-like structure hinders the diffusion of particles through it. Any aerosolized treatment of lung diseases must have certain properties to circumvent the mucosal barrier, and these properties may vary between diseases, drugs, and patients. Using computational fluid dynamics, a model of this mucus layer was constructed to simulate the convective and diffusive transport of impacted aerosol particles. The model predicts the dosage fraction of particles of a certain size that penetrate the mucus and reach the underlying tissue, as well as the distance downstream of the dosage site where epithelial concentration is maximized. Reactions that may occur in solution are also considered, with simulated data for the interaction of a model virus and antibody. The model is modular so that various lung regions and patient health states may be simulated.

## 1. Introduction

Lung diseases afflict hundreds of millions of people and are some of the most common causes of death worldwide (World Health Organization, 2020). Existing methods of treating these diseases are limited by poor targeting to the lungs when medications are given orally or intravenously and require rigorous, long-term, often invasive medication to achieve remission in pathogenic diseases (Feng et al., 2018; Kolewe et al., 2021). Other diseases are chronic but suffer similar limitations in treatment methods, inhaled or otherwise. The idea of treating lung diseases at the source is an attractive one, offering a noninvasive route to locally dose a diseased area that also minimizes side effects; however, several problems still exist (Duncan et al., 2018; Hastedt et al., 2016; Henning et al., 2010). A significant physical challenge to overcome is the mucociliary clearance mechanism (Fig. 1), which utilizes rhythmically waving ciliated cells to constantly push the mucus layer that coats the lungs upward towards the throat. This system serves as a natural defense against infection and particle buildup but simultaneously acts as a barrier to drug delivery (Carlson et al., 2018; Chen et al., 2019; Duncan et al., 2016).

**Figure 1:**
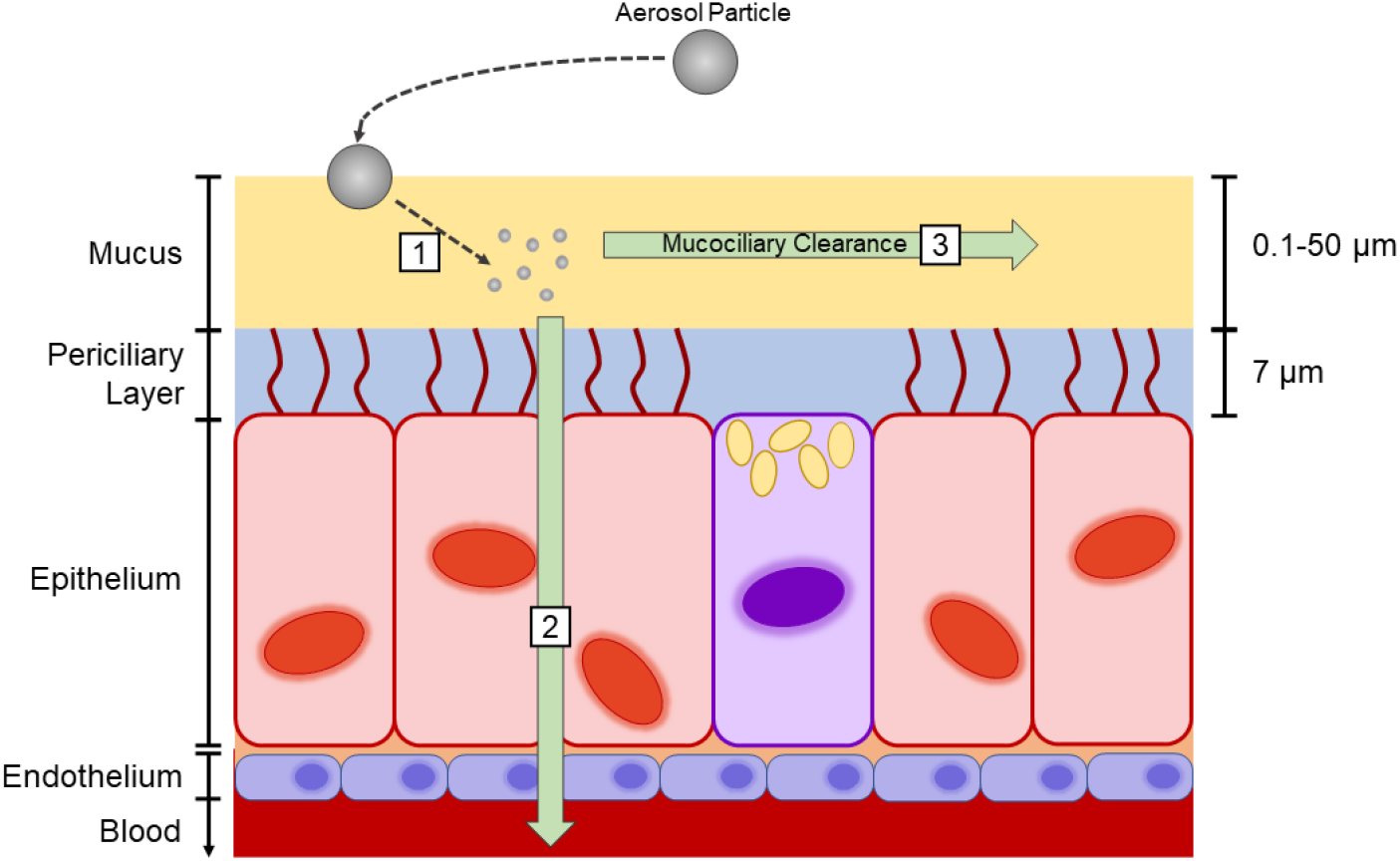
Methods of particle clearance from the lung mucosa. (1) Particles can be lodged in the mucus, where they may be degraded or consumed by macrophages before subsequent mucociliary clearance. (2) Particles can diffuse across the mucosa and be taken up into the tissue or even the bloodstream. (3) Most commonly, particles are either unable to diffuse adequately or are immobilized by some means and cleared by mucociliary clearance.

The anatomical interface between the airways and the lung epithelium is comprised of a fluid bilayer, with the highly viscous mucus layer on top of the more watery periciliary layer (PCL) (Fahy and Dickey, 2010; Taherali et al., 2018) (Fig. 1). The PCL allows the cilia to beat without catching in the mucus and acts as a lubricant that allows the mucus to slide along the interface (Cone, 2005; Chateau et al., 2018). The mucus is primarily water, but a network of glycoproteins known as mucins causes it to move as a bulk “sheet” when propelled by the cilia. Mucins are rich in cysteine, making them largely anionic (Cone, 2005). Disulfide bonds are a large contributor to the structure of the network. However, large portions of the chains are neutral, making mucins also appreciably lipophilic (Murgia et al., 2018). The PCL also contains mucins, but these are tethered to cilia and do not form the tangled net that is seen in the mucus layer. This makes the PCL a “brush” that prevents mixing of the two layers and maintains the viscous character of the PCL compared to the more elastic character of the mucus layer (Button et al., 2012). Because the cilia are regular in size, the PCL is consistently ≈ 7 µm thick through the entire tracheobronchial tree (Fahy and Dickey, 2010). In contrast, the mucus layer thickness varies significantly depending on location (Fig. 2). Goblet cells, which secrete mucus, are present throughout the respiratory tract and are more plentiful in the upper airways (Clarke and Pavia, 1980). Thus, the mucus layer increases in thickness from the lower airways to the upper (Fig. 2). Mucus can be as thin as 0.1 µm at the alveolar level (at the deepest levels of the respiratory tract) and as thick as 100 µm at the trachea (Chateau et al., 2018; Taherali et al., 2018). The velocity at which mucus is cleared depends on a variety of factors that vary significantly between individuals including age, health, and history of smoking.

**Figure 2:**
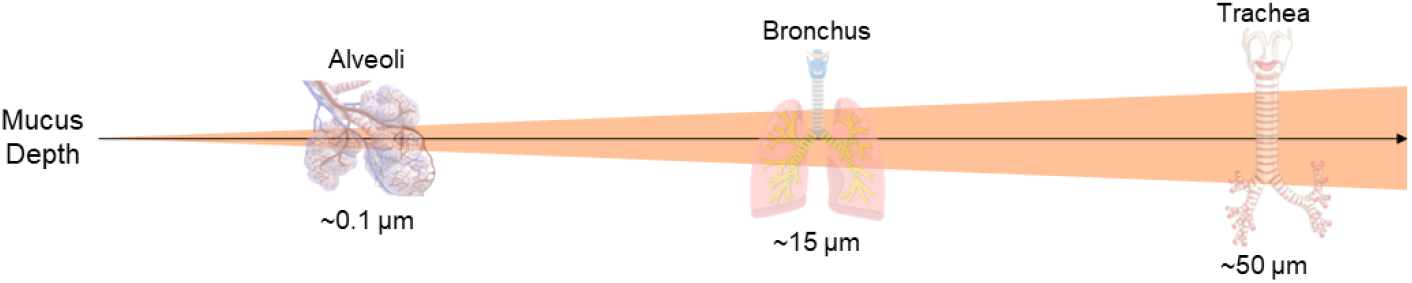
Mucus layer thickness thins toward more distal regions of the lung. Values given are estimates for healthy individuals. Note: background wedge indicating size variations is not to scale.

The tangled network of mucins in the mucus layer gives the layer marked elastic properties so that it behaves as a viscoelastic gel, which for our purposes is approximated as a shear-thinning non-Newtonian fluid (Lai et al., 2007; Cone, 2008; Norton et al., 2011; Murgia et al., 2018). This shear-thinning relationship exists in healthy individuals (and is the reason coughing is an effective method of clearing mucus (Taherali et al., 2018; King et al., 1985)) but is accentuated in obstructive lung diseases like cystic fibrosis, chronic obstructive pulmonary disease (COPD), and asthma. In these diseases, mucin production is elevated, causing the mucus to become much thicker and more difficult to clear than in the healthy case (Duncan et al., 2016). This “thickening” of the mucus, along with other factors like dysfunction in electrolyte concentrations, can lead to a collapse of the PCL as one or both layers become too viscous for the cilia to beat effectively (Taherali et al., 2018). Using a non-Newtonian mathematical model for the mucus allows for a single simulation that can be applied to both healthy and diseased states at the discretion of the user without extensive viscosity data, which is particularly lacking for diseased states.

The rheological properties of lung mucus must be taken into account to successfully treat lung disease via an inhaled aerosol, and the drug particles must have certain properties to pass through the mucus quickly and efficiently. A model of these fluid layers can predict the effective penetration of a delivered pharmaceutical or can be modified to simulate the infectivity of a pathogen. While researchers have been optimistic about the possibility of aerosol treatments for various lung conditions and diseases for decades (Yeates et al., 1975; Wolff, 1986; Lethem, 1993), poor characterization of the mucosa has been a limitation to progress. Experimental and theoretical developments in the understanding of the production of nanoparticles, aerosolization, and delivery to specific sites in the lung have made these treatments much more attainable (Tang et al., 2009; Kleinstreur and Feng, 2013; Feng et al., 2019), but the literature lacks a generalized model for predicting behavior in the mucus. Models that exist largely focus on cilial beating (Smith et al., 2008; Norton et al., 2011) or individual pores of the mucus layer (Cu and Saltzman, 2009; Hansing and Netz, 2018a,b). Other models are simplified to the point of ignoring fluid movement or the existence of multiple fluid layers (Kirch et al., 2012; Sims et al., 2019)–both of which leave questions about locating dosage sites or macroscopic behavior unanswered. The purpose of this research is to help fill that gap by producing a model that simulates the lung airway mucosal layers (Fig. 3) for a region of the tracheobronchial tree and that predicts the behavior of applied aerosolized particles. In this paper, we construct a computational fluid dynamics model of the lung airway mucosa informed by properties taken from the literature. After defining the model in Section 2, we show model predictions of the dosage distribution of particles of different sizes that penetrate mucus of various thicknesses to reach the underlying tissue, and we show the distance downstream of the dosage site where epithelial concentration is maximized for various cases. We also consider reactions to account for interactions between a model virus and antibody delivered to the mucus.

**Figure 3:**
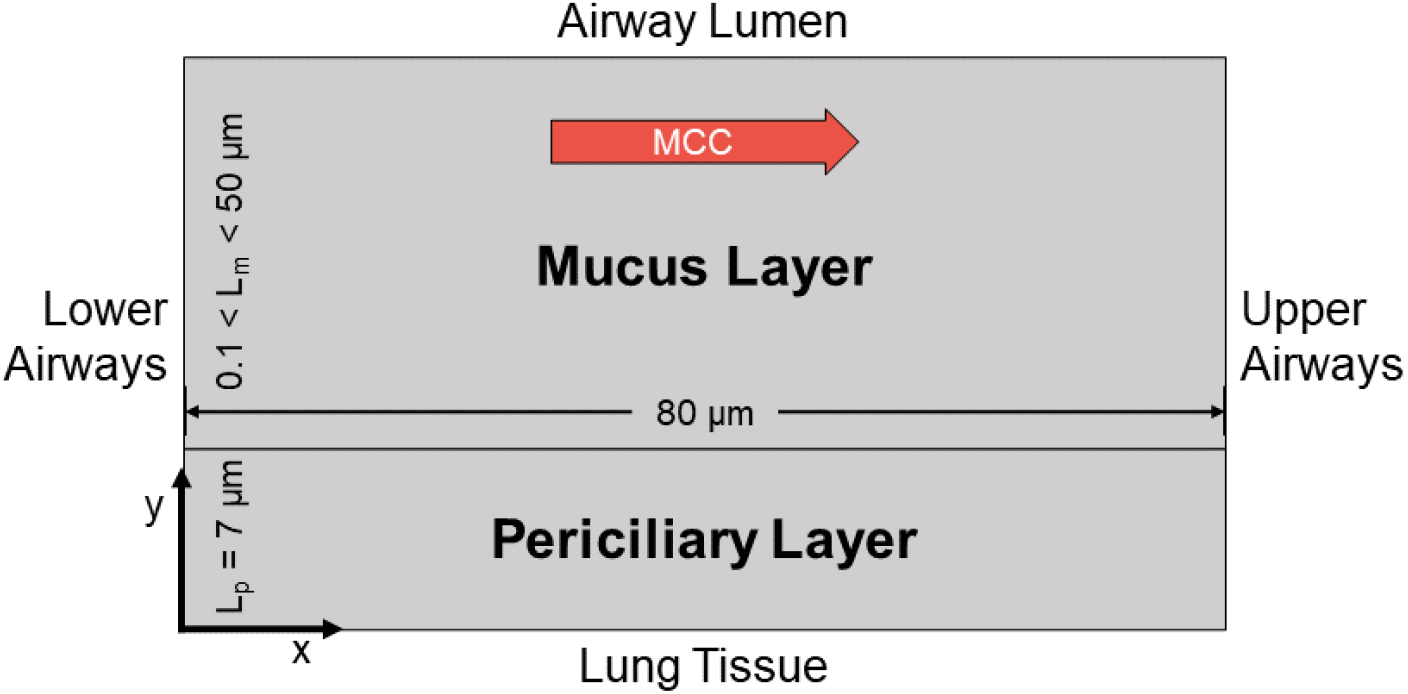
Example of the geometry used in the model. The top layer is the mucus and is open to the conducting airways, also called the lumen. The bottom layer is the periciliary layer; the cilia beat through this layer and are attached to cells of the lung epithelium. Mucus flows from the lower airways to the upper airways and the throat, and this directionality of the mucociliary clearance (MCC) is simulated as left to right in the diagram. *L*_*m*_ is the thickness of the mucus layer, which varies by region, and *L*_*p*_ is the thickness of the periciliary layer, which is uniform.

## 2. Methods

### 2.1. Model Geometry, Mesh, and Boundary Conditions

We used COMSOL Multiphysics 5.6 for modeling convection and diffusion through the mucus and PCL. The model geometry represents a cross-section of the mucosa at any particular location in the tracheobronchial tree. We assumed that the tracheobronchial tree is radially symmetric; thus, we consider a 2D rectilinear domain (Fig. 3). The domain does not need to be expanded to the left of the dosage site as fluid convective velocity is several orders of magnitude higher than diffusive velocity; therefore, backflow is negligible. The thickness of the simulated domain is dependent on the lung location being simulated (Fig. 2) but is always on the scale of microns, so all geometrical dimensions for the model are on this scale. The thickness of the periciliary (lower) layer is effectively constant throughout the tracheobronchial tree at 7 µm. The length of the tracheobronchial tree is on the scale of decimeters (Chateau et al., 2018). While the width of the model domain is set small enough to assume no significant changes in mucus thickness over the simulation domain, the specific domain width value is partially arbitrary. All simulation domains shown in this paper have a total width of 80 µm, as this value is sufficiently large to observe epithelial concentration maxima for all studied particle sizes.

The simulation domain of the model is two-dimensional, consisting of a series of four rectangles, arranged such that both a dosage site and a “downstream” area can be simulated with two distinct fluid layers (Fig. 4). Where the edges of these rectangles meet, COMSOL forms a union between them, such that there is no formal boundary condition separating them and all meshing is continuous. The top two rectangles represent the mucus layer, and the bottom two rectangles represent the underlying PCL. The rectangles on the left have the same fluid properties as the rectangles on the right with respect to their layer, but the left side functions as the dosage site, whereas the right side represents the fluid upstream of this site. For these simulations, we assume that concentration at the top edge of the dosage site is uniform to simulate behavior averaged across inhalation patterns, radial distribution, and other random transient effects related to the delivery of the aerosol itself, although an *in vivo* case likely follows an orientation-dependent distribution curve (Feng et al., 2018; Kolewe et al., 2021).

**Figure 4:**
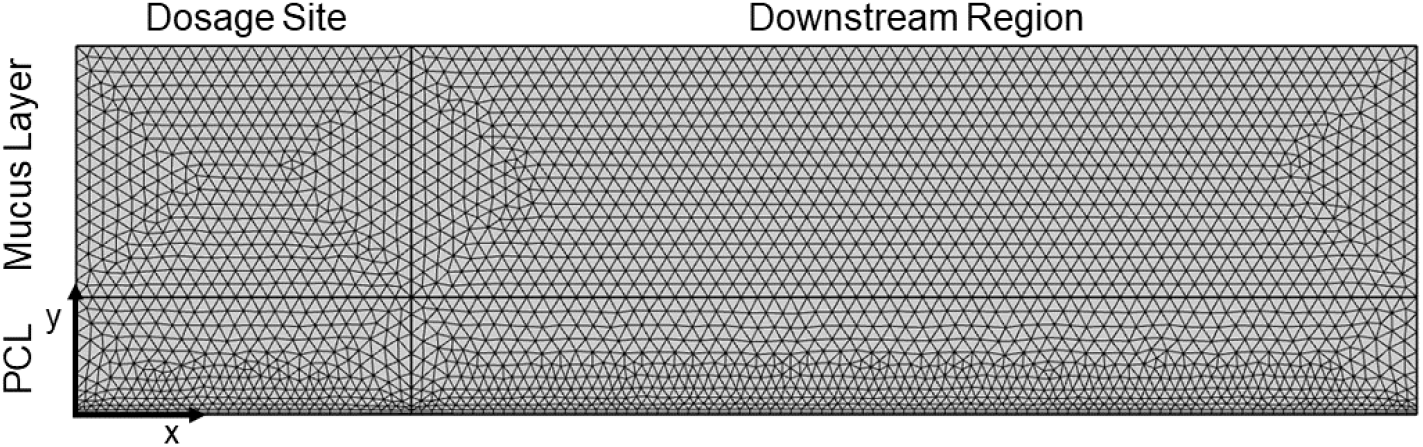
Mesh for the simulation domain. Four rectangles compose the simulation domain. In clockwise order from the top left, these represent the dosage region of the mucus layer (where the dosage is applied on the top surface at the air-mucus interface), the downstream region of the mucus layer, the downstream region of the periciliary layer (PCL), and the dosage region of the periciliary layer.

We use COMSOL Multiphysics to generate a mesh for the model using the “fine” resolution setting following a mesh independence test. This triangular mesh provides discrete points at which each relevant system of equations is solved, resulting in a two-dimensional field of behavior across the simulation domain. The mesh may be refined manually at the price of increased computation times.

The upper boundary of the mucus is open to the airways, making it a free surface under general conditions, and is governed by a slip condition. Conversely, the lower boundary of the PCL borders the static epithelium and is subject to a no slip condition. Bulk fluid moves from the left boundary of both layers to an outlet at the right boundaries without accumulation within the simulation domain. The mucus and PCL average velocities are specified (Table 1) and held constant. High viscosity and low velocity mean that the mucus has a small Reynolds number, moving in the laminar flow regime. Drug particles (molecules, aerosolized liquid droplets, or dry nanoparticles) are treated as continuous dilute species for modeling purposes. For the simulations shown here, the dilute species enters from the airways at a constant concentration constraint applied to the air-mucus interface (top surface of the top left rectangle in Fig. 4), which makes steady-state simulations possible. The dilute species may exit at the rightmost boundary—simulating mucociliary clearance —or at the bottom boundary—simulating uptake into the epithelium, which is assumed to occur instantaneously when the solute reaches the bottom boundary. This behavior is accomplished using outflow boundary conditions on these edges. The dilute species cannot cross the upper boundary, as this would imply that the applied particles may freely vaporize. Nothing prevents the dilute species from crossing the leftmost boundary as it is given an outflow boundary condition. In practice backflow does not occur due to the differences in magnitude between convection and diffusion. At the initial condition, no drug is in the mucus or PCL.

**Table 1:** Model parameters. PCL: periciliary layer

| Parameter | Definition | Value | Units | Ref. |
| --- | --- | --- | --- | --- |
| $\rho$ | Density of fluid | 1000 | kg/m<sup>3</sup> | (Norton et al., 2011) |
| $T$ | Body temperature | 310.15 | K | - |
| $\mu$ | Viscosity of water at $T$ | $6.922 \times 10^{-4}$ | Pa · s | (Korson et al., 1969) |
| $u_{muc}$ | Mucus average velocity | 5 | mm/min in $x$-direction | (Shete et al., 2014; Taherali et al., 2018) |
| $u_{pcl}$ | PCL average velocity | 2.4 | mm/min in $x$-direction | (Matsui et al., 1998) |
| $C_{Ai}$ | Dosage concentration | 1000 | mol/m<sup>3</sup> | - |
| $L_m$ | thickness of mucus | 0.5 – 100 | μm | (Chateau et al., 2018; Taherali et al., 2018) |
| $L_p$ | thickness of PCL | 7 | μm | (Fahy and Dickey, 2010) |
| $\phi$ | Fiber volume fraction | 0.0025 | unitless | (Hansing and Netz, 2018b) |
| $r_f$ | Mucin fiber radius | 5 | nm | (Lai et al., 2009) |
| $\mu_0$ | Mucus zero-shear-rate viscosity | 302 | Pa · s | Fitted to data (Lai et al., 2007) |
| $\sigma$ | Mucus shear thinning parameter | 8440 | s | Fitted to data (Lai et al., 2007) |
| $n$ | Mucus Carreau model exponent | 0.375 | s | Fitted to data (Lai et al., 2007) |

With the dilute species inlet being a constant concentration constraint and instantaneous outflow boundary conditions (no accumulation of the dilute species within the domain), the model domain reaches a steady state after a “startup time” has elapsed. For a domain of this size, this time is about 1 second. This makes a time-independent study possible. Such simulations are useful as they take a comparatively short time to compute and show long-term trends, most notably the concentration profiles.

### 2.2. Model Equations and Parameters

In this study, mucus and PCL flows are considered laminar and incompressible. Nanoparticles are modeled as a continuous dilute species. The nanoparticle-laden mucosal flows are simulated by solving the following governing equations, including the conservation laws of mass and momentum. The convection-diffusion equation (i.e., material balance equation) for the concentrations of particles and the possible products generated by reactions occurring within the mucosa is

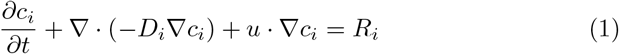

where *c*_*i*_ is the concentration of species *i, D*_*i*_ is diffusivity of species *i*, and *u* is fluid velocity. The first term represents accumulation in the domain, the second for diffusive transport, the third for convective transport, and the right side of the equation for the production or consumption of species as a result of chemical reactions. These equations require a value for diffusivity to be specified, which is defined later in this manuscript in (3). In the case where a chemical reaction is occurring to produce or consume the relevant species, the term *R*_*i*_ is defined as the net production rate of species *i*. If no reaction occurs, then *R*_*i*_ = 0. Other assumptions include constant density for the two layers (Norton et al., 2011), constant velocities for the clearance of the mucus layer and PCL (Matsui et al., 1998; Shete et al., 2014; Taherali et al., 2018), and equivalent diffusivity through the different layers. The fiber volume fraction for a healthy individual is in the range of 0.0005 to 0.01 (Hansing and Netz, 2018b). For all simulations in this manuscript, the fiber volume fraction was set to 0.0025. Mucin dimensions are within ranges reported by (Lai et al., 2009). Table 1 includes the values or ranges of the parameters used for the simulations.

For the diffusivity of spherical particles in bulk solution, we use the Stokes-Einstein equation

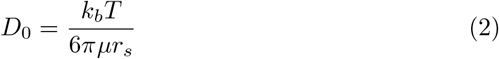

where *k*_*b*_ is the Boltzmann constant, *T* is absolute temperature, *µ* is the dynamic viscosity of the pure solvent (in this case water), and *r*_*s*_ is the Stokes radius of the solute particle. We allow *r*_*s*_ to vary throughout our simulations. However, mucins provide a steric hindrance to diffusion, which we account for by using an appropriate effective diffusivity correlation (3). The mucus is porous with an average pore size of around 150 nm (Button et al., 2012; Kim et al., 2016), but the network is not rigid. Mucus is reasonably described as a hydrogel, where diffusivity is a function of fiber size rather than a function of pore size. We solve for an effective diffusivity, *D*, for a solute moving through the mucus and PCL using a correlation from the literature that accounts for both steric and hydrodynamic interactions in a fibrous hydrogel (Philips, 2000):

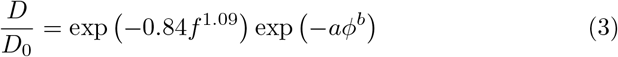

where *λ* = *r*_*f*_ */r*_*s*_, *r*_*f*_ is the fiber radius, *r*_*s*_ is the solute particle radius, *ϕ* is the fiber volume fraction, and

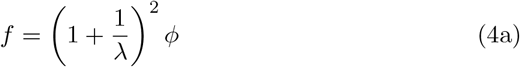

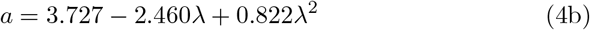

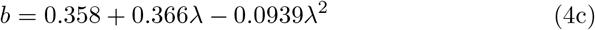

Equations (3) and (4) are fully characterized by only three parameters: fiber radius (*r*_*f*_), particle radius (*r*_*s*_), and fiber volume fraction (*ϕ*). Particle radius is one of the primary design variables for this study. No specific pore size or shape is assumed in (3). Rather, hindrance is related to the likelihood that a diffusing particle will collide with a fiber. Additionally, (3) does not rely on Brinkman or effective medium approximations. The Brinkman equation is a variation of Darcy’s law that is designed to describe flow in media where the grains of the media are themselves porous and requires measurement of effective viscosity. Effective medium theory as applied to these situations, in short, considers the mucus to be characterized only by its Darcy permeability (Johnson et al., 1996). Both effective viscosity and Darcy permeability are generally more difficult to calculate, measure, or estimate than *r*_*s*_, *r*_*f*_, and *ϕ*. Thus, (3) is more approachable than similar equations that rely on these approximations, and (3) also tends to fit data more accurately (Philips, 2000). The form of the conservation of momentum equation used by the COMSOL Multiphysics software is

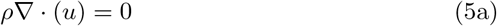

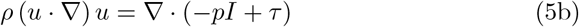

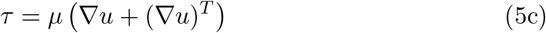

where *ρ* is the fluid density, *p* is fluid pressure, *u* is the fluid velocity, *µ* is the fluid viscosity, and *I* and *τ* denote the identity tensor and the stress constitutive equation, respectively. Reported measurements of clearance rates were collected *in vivo* and thus already include the effects of gravity and breathing. Consequently, the gravity term is neglected from the momentum equation to avoid double counting of effects that were not decoupled in the experiments. Since the flow regime is open to the airways, the mucus moves by open channel flow, and flow is not pressure-driven. These equations constitute the Navier-Stokes equations for an incompressible fluid and are subject to boundary conditions as specified above.

Considering the relation of mucus viscosity to shear rate (Vélez-Cordero and Lauga, 2013), we model the mucus layer as a Carreau fluid (Shahsavari and McKinley, 2015). The Carreau model is used in other applications to simulate similar biological fluids with non-Newtonian characteristics like blood. The governing equation is

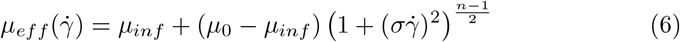

where *µ*_*eff*_ is used as *µ* for the mucus layer in (5c), 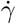 is the shear rate, *µ*_*inf*_ = 0 is the infinite shear-rate viscosity (assuming that viscosity is small at infinite shear so that this term drops out of the equation), *µ*_0_ is the zero-shear-rate viscosity, *σ* is the inverse of a characteristic shear rate at which shear thinning becomes important, and *n* is a power law exponent. Published experimental data was used to parameterize (6) for mucus. We used WebPlotDigitizer (Rohatgi, 2022) to digitize the data in (Lai et al., 2007). Then we used nonlinear least-squares regression to fit parameters *µ*_0_, *σ*, and *n* to the (Lai et al., 2007) data for cervicovaginal mucus. The values are listed in Table 1. The raw data was for cervicovaginal mucus. Cone (2008) showed that all sources of mucus have very similar shear-thinning viscous behavior except ovulatory cervicovaginal mucus. As the data in (Lai et al., 2007) is non-ovulatory, it was assumed to be a reasonable proxy for the viscous behavior of lung mucus.

### 2.3 Code Availability

We have provided the COMSOL code and exported files for the results in a repository at https://github.com/ashleefv/CFDparticleLungMucosa (Bartlett and Ford Versypt, 2023).

## 3. Results and Discussion

Before the diffusion of a dilute species was considered, we verified that the fluid velocity profile was realistic. In the biological system, the mucus layer is so viscous (due to the tangled network of mucins) that it moves largely as a single sheet. The PCL is more Newtonian due to conformational differences in mucin structure and is cleared more slowly because it does not respond elastically to cilial beating (Norton et al., 2011; Matsui et al., 1998; Hussong et al., 2013). Due to these differences in behavior, the Carreau correlation described in (6) was only applied to the mucus layer, and the PCL layer was treated as Newtonian with the viscosity of the PCL approximated as that for water. Average velocity *u*_*muc*_ in the *x*-direction was specified as 5 mm/min, an average *in vivo* tracheal mucus clearance rate (Shete et al., 2014; Taherali et al., 2018). Average velocity *u*_*pcl*_ was specified as 2.4 mm/min (Matsui et al., 1998). Fig. 5 shows the bulk movement of the mucus layer and the rapid drop in velocity in the PCL due to both the no-slip condition with the epithelium and property differences between the two layers.

**Figure 5:**
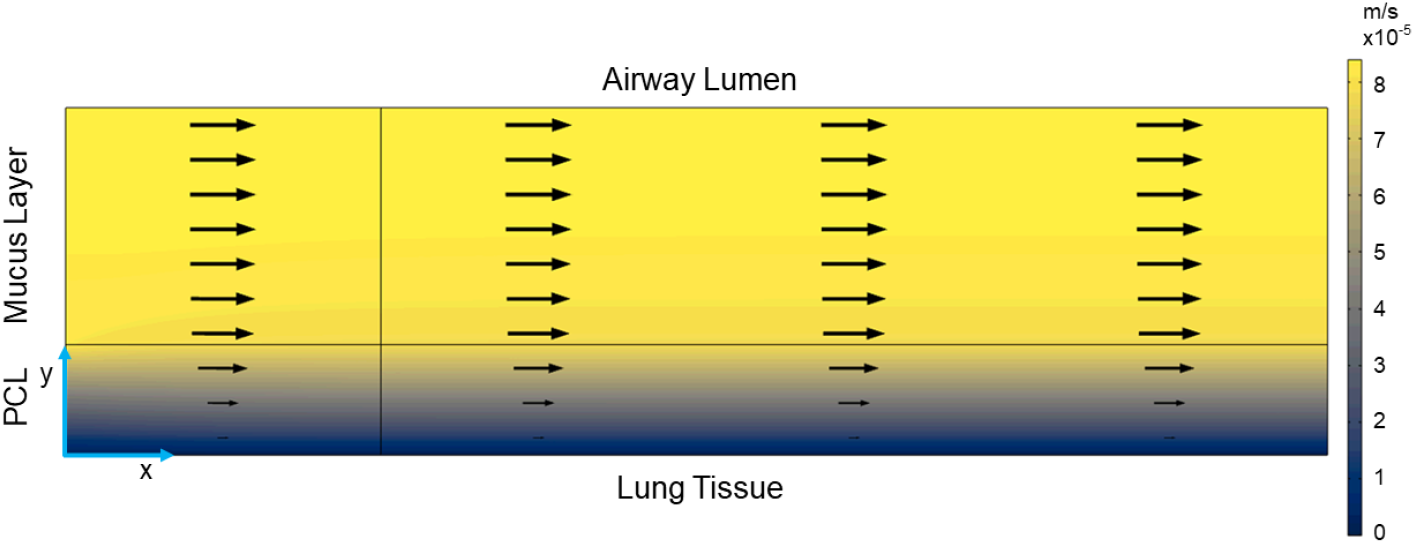
Velocity profile for *L*_*m*_ = 15 µm. Arrows point in the direction of mucus flow and scale in size with the velocity. Note a nearly constant velocity throughout the mucus (upper) layer.

Concurrent with the velocity profile arising from the viscous nature of the two fluids, it follows that the PCL is subject to a high shear rate (Fig. 6)), as cilia beat through it constantly. Conversely, the mucus layer is subject to a low shear rate due to its elasticity (Fig. 6). In short, the shear rate is another measure of the mucus moving as a sheet, as the bulk mucus moves in response to cilial beating. The PCL is also responsive to this beating, but to a much smaller extent due to both the no-slip condition for the velocity profile and the much lower viscosity of the PCL. Another way of interpreting Fig. 6 is as a measure of resistance to movement. The high shear rate in the PCL is indicative of its Newtonian character, where the lack of elasticity results in a high shear rate required to achieve the observed velocity profile. The cause of this profile *in vivo* is cilial beating, but defining the velocity profile results in identical bulk fluid mechanics so cilia do not need to be explicitly modeled. Similar to the velocity profile, the values are nearly constant in the mucus but subject to a gradient in the PCL, albeit with high values in the PCL instead of the mucus in contrast to relative values of the corresponding layers in the velocity profile.

**Figure 6:**
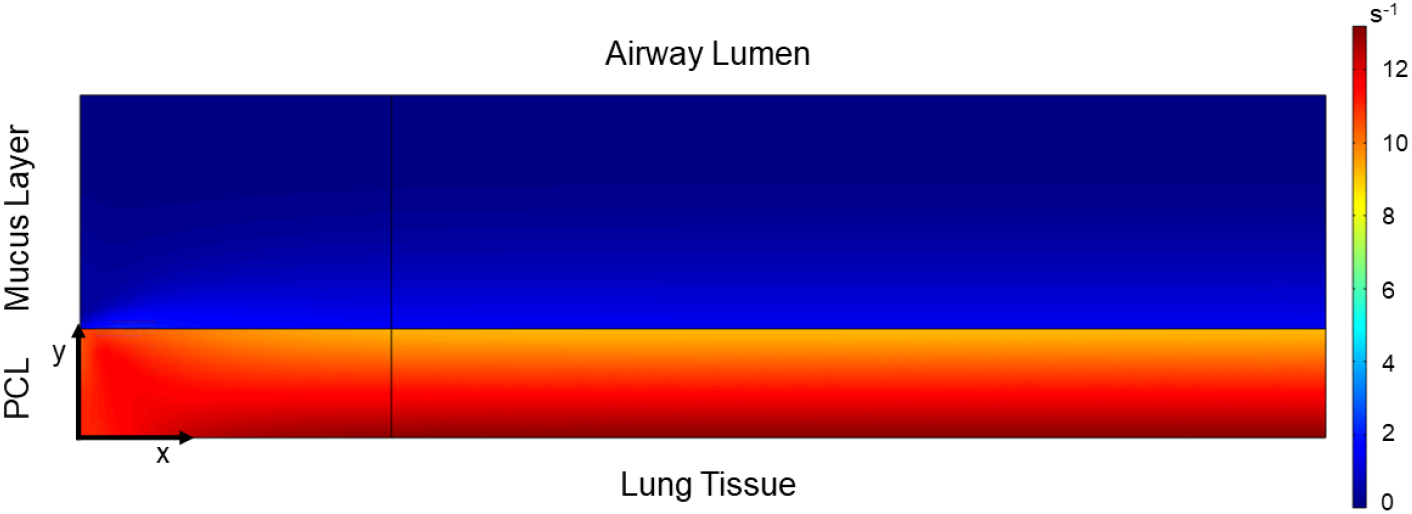
Shear rate in the mucus (upper) and periciliary (lower) layers. The mucus (upper) layer has a comparatively small and constant shear rate due to its elasticity.

Once a velocity field was constructed, a drug dilute species was applied to study its transport through the domain. All simulations in this paper used a uniform delivery concentration of 1000 mol/m^3^ = 1 mol/L along the lumen-side boundary of the dosage site. In the delivery of an actual drug, delivery would likely not be constant. Here, all mucus within the domain had a residence time of about 1 second due to mucociliary clearance, so this was considered a short enough timescale for constant delivery to be valid. It was assumed that this concentration is sufficiently dilute that the delivered drug does not significantly change the volume of the system. COMSOL supports parameter sweeps, where a single solution of the system evaluates a range of parameters. This functionality was first utilized to compare the effect of mucus layer thickness (*L*_*m*_) on drug penetration. Concentration profiles for three different mucus thicknesses, *L*_*m*_ = 5, 15, and 33 µm, were obtained (Fig. 7) to represent delivery to the bronchioles, bronchus, and trachea, respectively. Drug particle radius was held constant at *r*_*s*_ = 20 nm.

**Figure 7:**
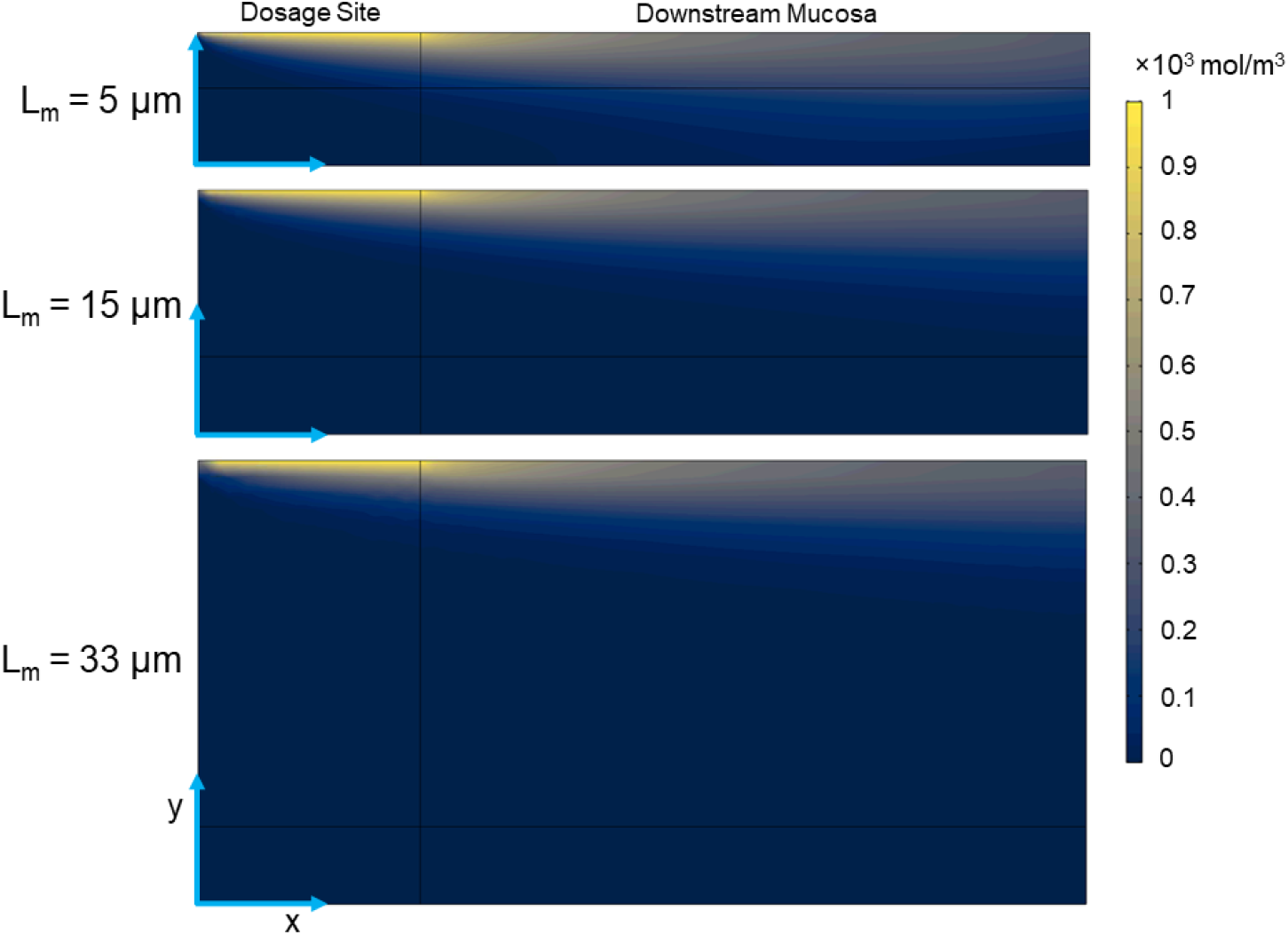
Drug concentration profiles for particles with a radius *r*_*s*_ = 20 nm for various mucus thicknesses (from top to bottom): *L*_*m*_ = 5, 15, and 33 µm. The applied concentration constraint at the dosage site is 1000 mol*/*m^3^.

Parameter sweeps for drug particle radius were also performed (Fig. 8). Mucus thickness was held constant at *L*_*m*_ = 15 µm. Concentration profiles for particles of radii *r*_*s*_ = 5, 20, and 60 nm are shown, highlighting the strong dependence of particle size on effective diffusivity. These sizes were chosen arbitrarily but are in the range of many viruses and engineered nanoparticles (Fig. 9). The profile for the *r*_*s*_ = 20 nm particle under these conditions is identical to the middle profile in Fig. 7. Larger particles are much more susceptible to steric hindrance in the mucus. While other factors are also at play, for many pathogens the basic reproduction number (*R*_0_) seems to trend inversely with size, with large pathogens like tuberculosis being generally less infective than smaller pathogens like ebolavirus (Delamater et al., 2019) and SARS-CoV-2.

**Figure 8:**
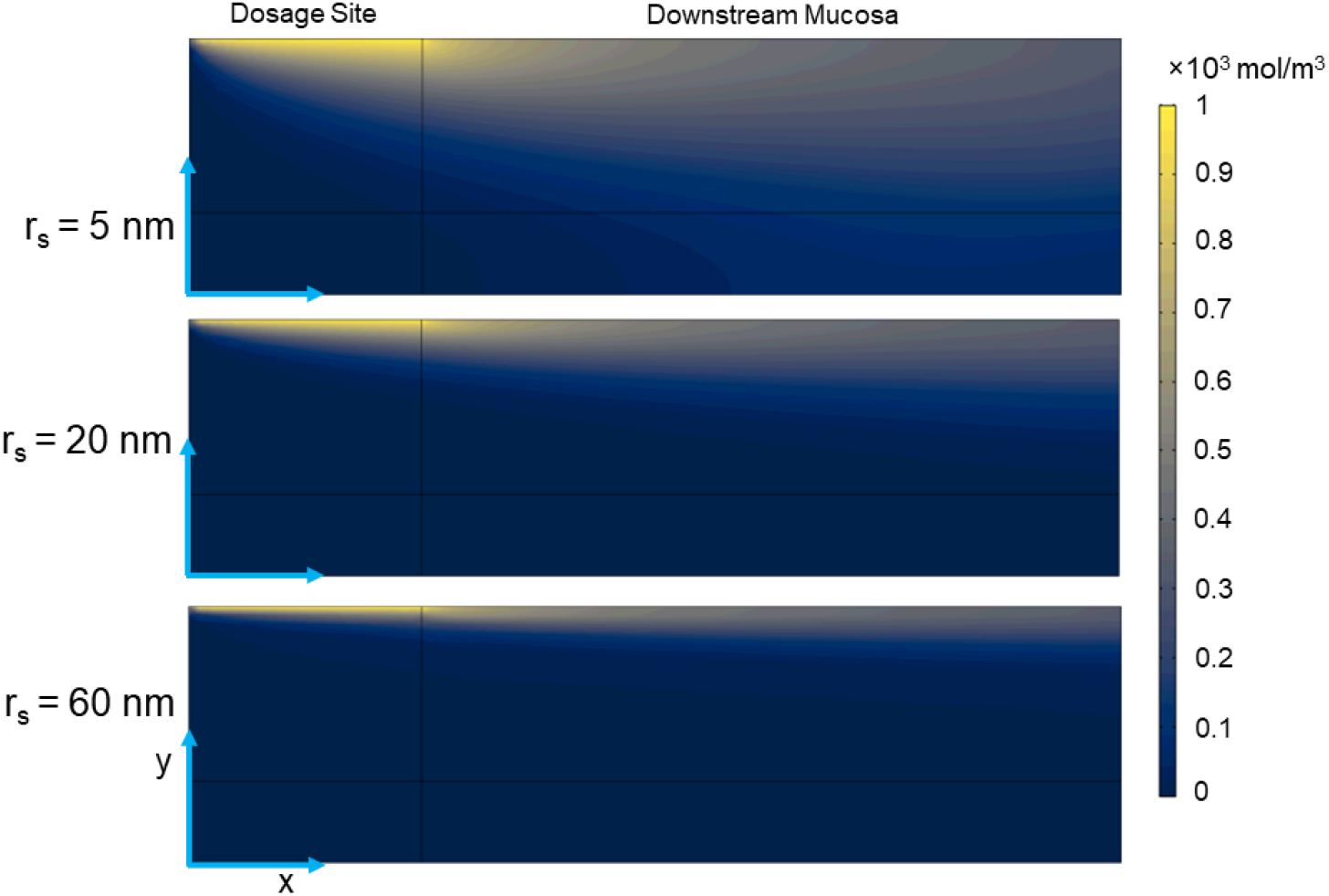
Drug concentration profiles for a mucus thickness of *L*_*m*_ = 15 µm and particles with radii (from top to bottom): *r*_*s*_ = 5 nm, 20 nm, and 60 nm. The applied concentration constraint at the dosage site is 1000 mol*/*m^3^.

**Figure 9:**
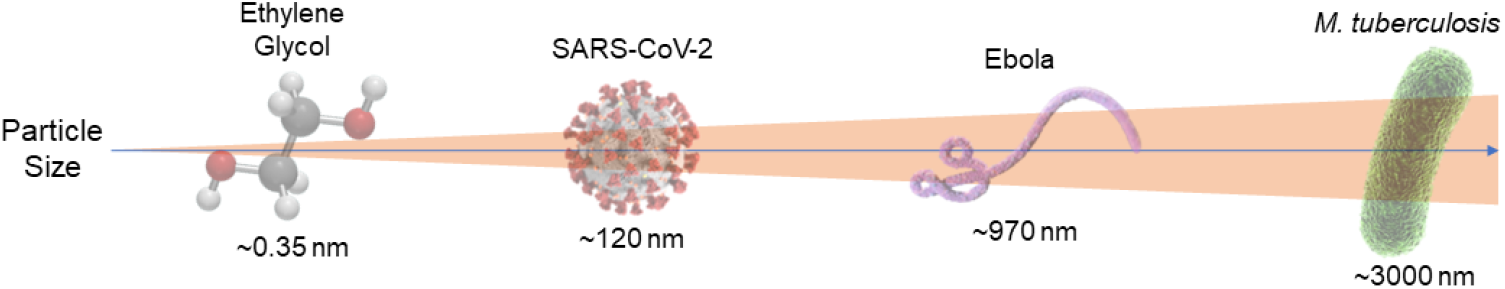
Examples of molecules and pathogens on the size scale used in these simulations. Ethylene glycol was selected as a representative molecule because of the prevalence of using polymers consisting of ethylene glycol monomers for many drug delivery applications. Note: background wedge indicating size variations is not to scale.

As expected, thicker mucus layers are more effective barriers to particle transport than thinner layers, and larger particles have lower diffusivities than those of smaller particles. This is generally true for passive diffusion in any fluid but is particularly relevant here as drug that does not fully cross both fluid layers is ultimately removed from the lungs by mucociliary clearance and then eliminated. Particles deposited deeper in the lungs will have greater success penetrating the mucus and reaching tissue (higher bioavailability) than the same particles deposited less deeply in the lungs.

Maximizing dose depth into airways is not always the goal, as certain patients may benefit from a specific localized dosage or tissue targeting along the tracheobronchial tree, such as in cancer treatment (Kolewe et al., 2021). For treatment regimes such as these, it is important to note that an appreciable amount of the applied drug only reaches the epithelium some distance upstream of the dosage site. For a particle in any mucus thickness, there is some distance upstream where the delivered dosage is maximized. By adding a cut line at the lung tissue epithelial surface and exporting the concentration results along this *y* = 0 line, this distance was found for various particle radius and mucus thickness combinations (Figs. 10 and 11). The smallest particles reach the epithelium quite close to the inlet (dosage site) when *L*_*m*_ = 5 µm, whereas in thicker mucus they reach the epithelium further from the inlet and have maxima that are smaller in magnitude than the thinnest mucus case. These results show how far a targeted “dosage site” for the applied drug needs to be upstream of the epithelial position where drug concentration is maximized. These results also could be integrated to determine the entire area under the curve, a common dosage metric for the total amount of the dosage that reaches the target location. We used 10^*−*12^ mol/m^3^ as the minimum threshold concentration in Fig. 11 to ensure that the concentration that reaches the tissue is sufficiently large to be effective. Another important trend to notice is that both increased particle size and increased mucus thickness result in “flattening” of the curves shown in Figs. 10 and 11. Increases to either of these parameters result in the maxima occurring further downstream and being smaller in magnitude. When a treatment is designed, the simulation results for those drug particles can be used to calculate a dosage and delivery scheme. A dosage can be selected such that the desired uptake concentration to achieve a physiological response is achieved at the *x*-position corresponding to the maximum concentration (or a concentration above an acceptable threshold), and a dosage site selected based on the upstream distance required for this peak to occur at the target site. Alternatively the area under the curve of the concentration that reaches the epithelium could be maximized for a given zone.

**Figure 10:**
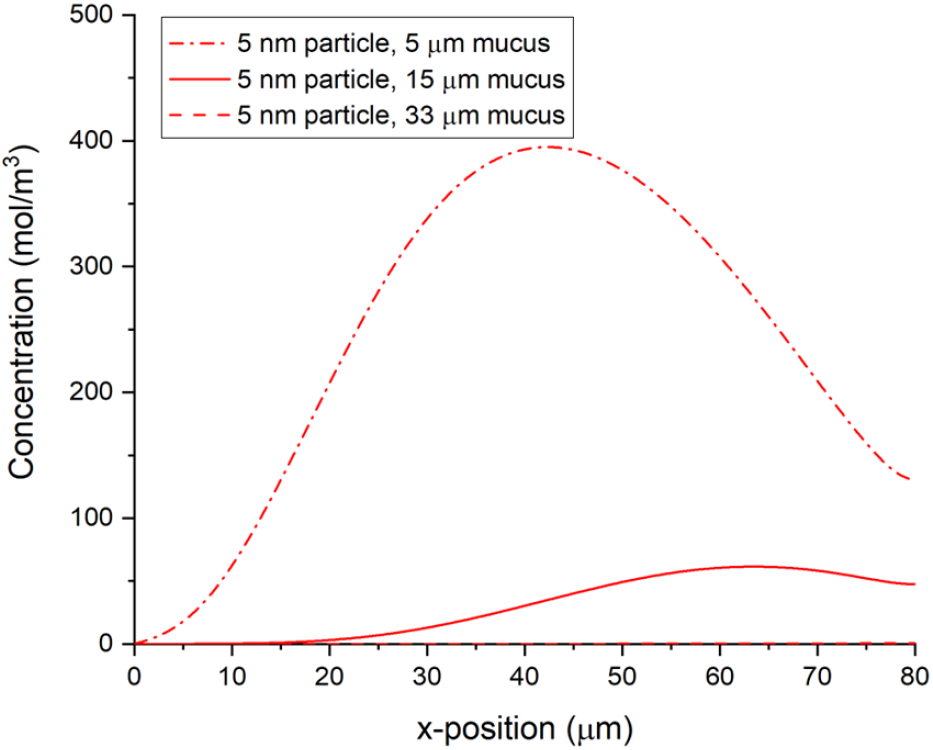
Drug concentration at the epithelial surface *y* = 0 as a function of *x*-position in the steady-state model for particles with r_s_ = 5 nm, where each curve has a different thickness of the mucus layer (*L*_*m*_) indicated in the legend. The applied concentration constraint at the dosage site is 1000 mol*/*m^3^.

**Figure 11:**
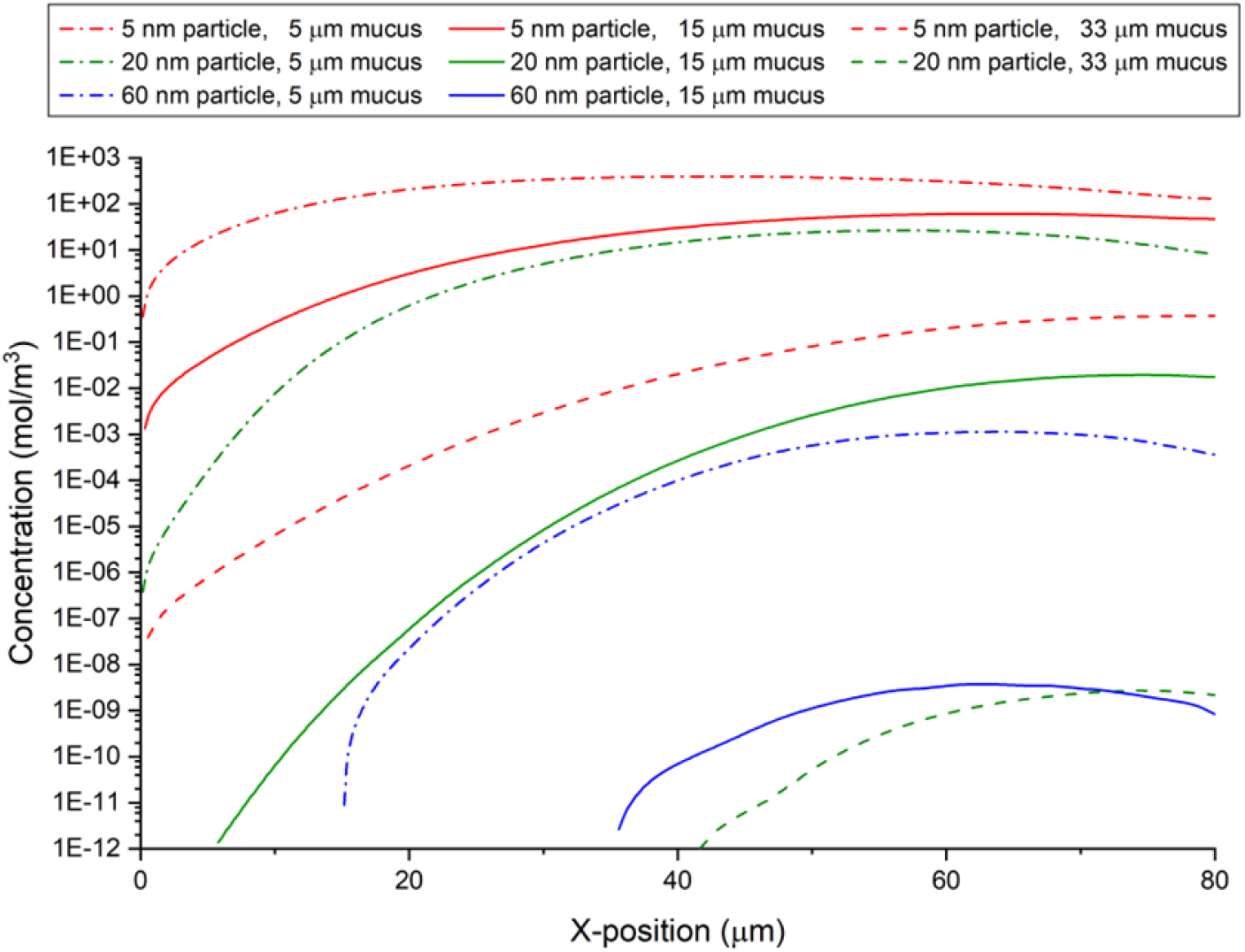
Drug concentration (log scale) at the epithelial surface *y* = 0 as a function of *x*-position in the steady-state model for combinations of three mucus thicknesses (*L*_*m*_) and particle sizes (*r*_*s*_), selected for having values on similar orders of magnitude to demonstrate the interplay between particle size and mucus thickness. The applied concentration constraint at the dosage site is 1000 mol*/*m^3^. Note that the case of *r*_*s*_ = 60 nm and *L*_*m*_ = 33 µm was also simulated, but the values are below the 10^*−*12^ mol/m^3^ concentration threshold.

The model provides information regarding the magnitude of dosage delivered along the lung epithelium. However, factors that may optimize delivery such as shrinking the particle size or delivering deeper into the lungs may not be viable. Particle size causes variations in impaction with the mucus when inhaled (Schlosser et al., 2010), and a particle may require a protective coating to prevent side reactions, denaturation, or electrostatic entrapment (Huang et al., 2017; Halwes et al., 2018; Osman et al., 2018; Patil et al., 2018). It has been hypothesized that drugs may be applied that slow the beating of cilia, thereby decreasing mucus clearance and giving the drug more time to cross the fluid layers. This is an interesting idea, but many pathogenic diseases of the lung are directly caused by mucociliary clearance dysfunction, so this method would likely increase the risk of complications (Nawroth et al., 2019). In non-pathogenic diseases characterized by mucociliary clearance dysfunction such as cystic fibrosis, asthma, and COPD, steric (and electrostatic) hindrance is often highly accentuated due to higher-than-normal mucin concentrations. In these diseases, additional drugs (or even simply water) might be applied to cause the mucus to behave more like the healthy case and improve outcomes (Nafee et al., 2018).

One major limitation of the model is the assumption that electrostatic interactions are negligible. This assumption is valid for many particles including many viruses and polyethylene glycol-coated nanoparticles but certainly not all particles (Cahn et al., 2023). Mucins are anionic and lipophilic, so cationic or lipophilic particles are subject to electrostatic hindrance, which can be immobilizing (Hansing and Netz, 2018b). Repulsive anionic interactions between particles and mucins result in a sort of channel flow, which is less hindering but still causes an effective shrinkage of pores. Particles that are surface neutral but hydrophilic (either polar or zwitterionic) typically are the most successful at penetrating mucus in the absence of any form of active transport.

Unwanted side reactions may be an issue in many aerosolized drug applications or side reactions could be desired in the case of a prodrug therapy. One specific application of interest is the simulation of prophylactics in the mucus. Lung mucosa, like other mucosal tissues, is an immunoactive region, with antibodies present in solution. Here, we considered a scenario when prophylactic antibodies against a virus particle have already been administered and are assumed to be uniformly distributed through the mucus in a zone. When these antibodies bind to an antigen (such as a virus), the resulting complex is often trapped sterically or electrostatically. We simulated this reaction as an elementary irreversible reaction between two species:

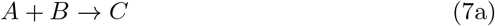

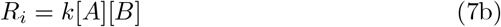

where species A is the inhaled particle (simulating a virus), species B represents a prophylactic antibody present in the mucosa, and product C represents an antibody-antigen complex. For these simulations, the rate law coefficient was defined as *k* = 0.05 m^3^*/*(mol · s). The applied concentration of A is *C*_*A,i*_ = 1000 mol*/*m^3^ at the dosage site (as in the other simulations), and the initial concentration of B is *C*_*B,i*_ = 20 mol*/*m^3^ present uniformly throughout the domain. Both concentrations were chosen arbitrarily; however, the radius of the viral particle A was set to *r*_*s,A*_ = 60 nm to simulate SARS-CoV-2 (Renu et al., 2020; Cascella et al., 2022). The radius of the particle B was set to *r*_*s,B*_ = 6 nm to simulate a monoclonal antibody (Hawe et al., 2011). We consider the complex C to be immobilized and thus has a diffusion coefficient of zero. The other two species have diffusion coefficients calculated using (3). Using these relationships, it can be shown whether a given level of antibody expression in the mucus is sufficient to prevent infection (defined as a certain amount of antigen reaching the epithelium) (Fig. 12). In these simulations, the diffusing particle is not a drug but a disease-causing viral particle. The model is equally capable of simulating the penetration of pathogens through the mucus as it is of therapeutic nanoparticles. Notice that some of species A persists at the rightmost boundary (Fig. 12) as all local concentration of B has been consumed. Species C has a notably different profile due to the immobilization of the species.

**Figure 12:**
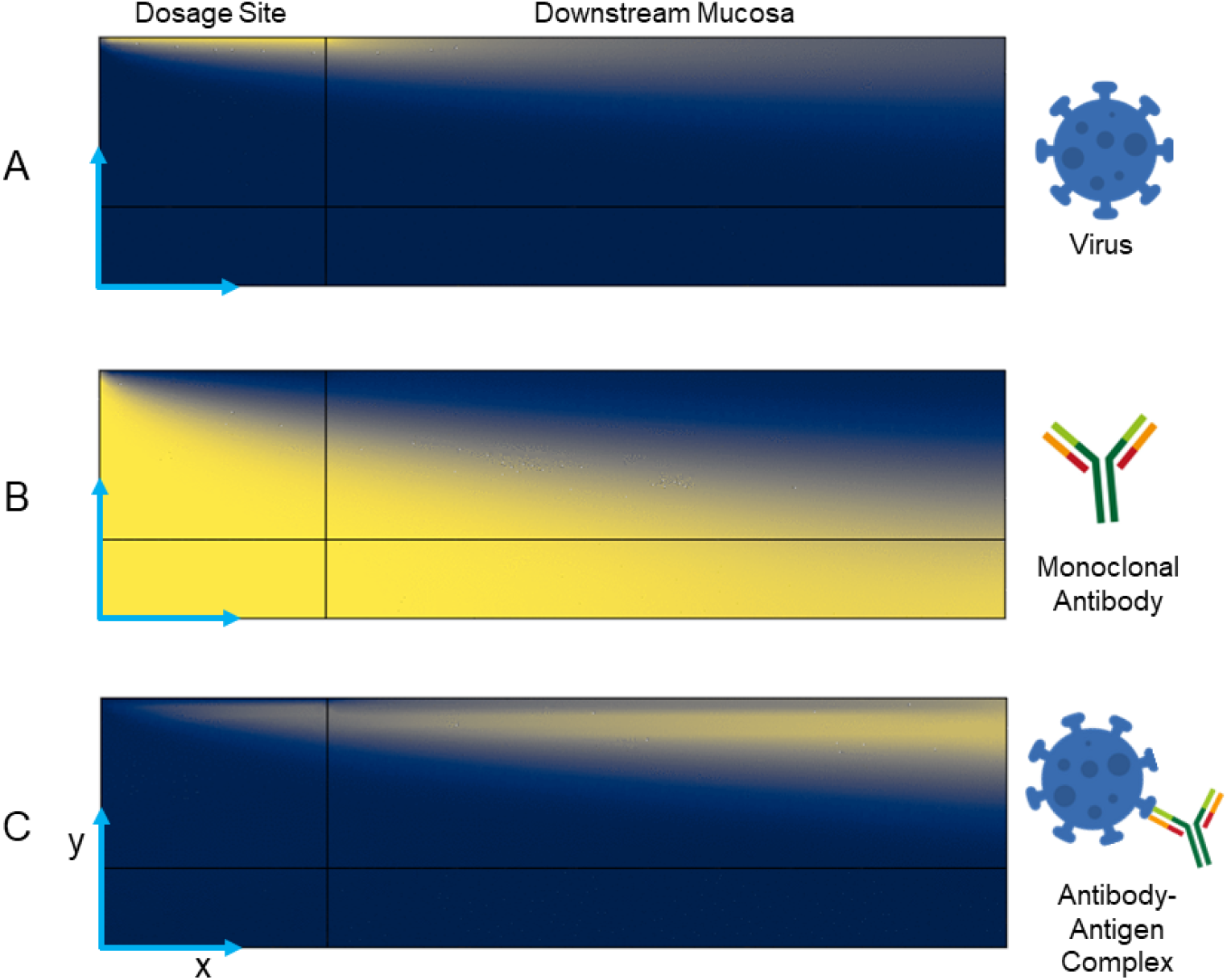
Concentration profiles of the three interacting species, where species A (virus) is uniformly applied at the dosage site and species B (monoclonal antibody) is initially present uniformly throughout the domain. Species A and B are consumed by the reaction, and species C is the non-diffusing product (antibody-antigen complex).

Using our model, users can solve for various unknowns when designing a drug delivery regimen. For example, for an existing drug particle of known size, a dosage site and dosage amount can be calculated via simulations such as those shown in Figs. 7 and 10. The efficacy of an administered prophylactic treatment may be screened via simulation of pathogenic exposure, such as that shown in Fig. 12, with the same approach being useful for simulation of prodrug therapies or controlled release of encapsulated drugs. The model can also predict whether a theoretical particle will significantly penetrate the mucus at all or be cleared from the body. The non-Newtonian physics of the mucus layer allows for the simulation of specific individual microenvironments or disease states by variation of only a few discrete variables that are simple to obtain experimentally. As many immunotherapies, especially those that are aerosolized, use spherical (or morphologically isotropic) particles, the model is well-suited to simulate these drugs, as relative diffusivities are calculated from the Stokes-Einstein equation. This means that morphologically anisotropic particles or pathogens, such as rod-shaped bacteria, spiral-shaped bacteria, and filamentous viruses will likely have behavior that deviates from the model. This is doubly true for pathogens that have evolved special mucopenetrative features, such as influenza type A (Vahey and Fletcher, 2019).

## 4. Conclusions

The model developed using COMSOL Multiphysics is a biologically realistic simulation of the mucociliary clearance mechanism. The model is customizable to the needs of the modeler or even the physiology of a patient, including both mucus properties and physical dimensions of the simulation domain. The administered particle is likewise customizable, although the model only accounts for steric and hydrodynamic hindrance. Existing simulations accounting for electrostatic interactions in “interacting gels” like mucus are computationally intense (Hansing and Netz, 2018a), and a macroscopic mathematical relationship that may be added as an additional term to (3) is needed. By exporting transport simulation results, plots of concentration reaching the epithelium versus distance downstream from the dosage site were created. Using this information the optimal dosage site was identified for some test cases. Such information has relevance in the development of aerosolized drug treatments for localized diseased tissues, including tumors. One possible future refinement is replacing the concentration inlet with a time- and position-dependent step function, which simplifies the domain to only an upper mucus layer and lower PCL and adds possibilities for non-steady-state simulations.

## 5. Acknowledgments

This work was supported by National Institutes of Health grant R35GM133763 (ANFV), Oklahoma State University: Freshman Research Scholar and Wentz Research Scholar Programs (BAB) and startup funds to ANFV, and the Gold-water Scholar Program (BAB). We would like to thank Ford Versypt lab members for their feedback on this manuscript.

